# Time-resolved multivariate pattern analysis of infant EEG data

**DOI:** 10.1101/2021.06.16.448720

**Authors:** Kira Ashton, Benjamin D. Zinszer, Radoslaw M. Cichy, Charles A. Nelson, Richard N. Aslin, Laurie Bayet

## Abstract

Time-resolved multivariate pattern analysis (MVPA), a popular technique for analyzing magneto- and electro-encephalography (M/EEG) neuroimaging data, quantifies the extent and time-course by which neural representations support the discrimination of relevant stimuli dimensions. As EEG is widely used for infant neuroimaging, time-resolved MVPA of infant EEG data is a particularly promising tool for infant cognitive neuroscience. MVPA methods have recently been applied to common infant imaging methods such as EEG and fNIRS. In this tutorial, we provide and describe code to implement time-resolved, within-subject MVPA with infant EEG data. A pipeline for time-resolved MVPA based on linear SVM classification is described and implemented with accompanying code in both Matlab and Python. Results from a test dataset indicated that in both infants and adults this method reliably produced above chance classification accuracy. Extensions of the core pipeline are presented including both geometric- and accuracy-based representational similarity analysis, implemented in Python. Common choices of implementation are presented and discussed. As the amount of artifact-free EEG data contributed by each participant is lower in studies of infants than in studies of children and adults, we also explore and discuss the impact of varying participant-level inclusion thresholds on resulting MVPA findings in these datasets.

## 1. Introduction

The contents of the infant mind are both fascinating and elusive. Without the benefit of verbal communication, inferring the mental states and representations of infants from behavior or neuroimaging data is an ongoing challenge. Functional imaging methods such as functional near-infrared spectroscopy (fNIRS) and electroencephalography (EEG) are popular in infant research due to their non-invasiveness and relative tolerance for movement while recording (Bell & Cuevas, 2012). These methods provide either fine-grained temporal with limited spatial information (EEG) or moderate spatial with limited temporal information (fNIRS) about neural responses, and typically consist of group average responses to stimuli (Dehaene-Lambertz & Spelke, 2015). While these methods can reveal information about conditional differences in timing or amplitude driven by different stimuli, they rely on averages from one or more electrodes/optodes (e.g., clusters of channels), ignoring information that may be represented in the patterns contained within these clusters.

Machine learning approaches including multivariate pattern analysis (MVPA) or “decoding” that have historically been used with adult neural data are promising avenues for infant research. Rather than finding differences in average stimulus-response recordings, MVPA is used to map patterns of activation across a cluster of channels to specific stimuli, or other relevant dimensions of the task or of the individual participant (Haynes & Rees, 2006). Using machine learning classification techniques, the goal of MVPA is to reliably discriminate between the patterns of activation associated with particular stimuli, categories of stimuli, or other relevant aspects of the participant’s phenotype (e.g., their attentional state or intrinsic trait). If patterns of neural activation can reliably map to stimuli (i.e., enable above-chance classification accuracy), it is plausible that these neural patterns support the discrimination of these stimuli, although we cannot infer whether the detected information drives behavior without manipulating these neural patterns (Haxby et al., 2014; Isik et al., 2014). This technique has been applied to adult data, primarily fMRI voxels, to index the information that can be extracted from brain activity, including in multivariate, spatially distributed representations (Haxby, 2012). Multivariate methods have been used in many research contexts and stimulus modalities including discrimination of painful stimuli (e.g. Brodersen et al., 2012), localized touch sensation (e.g. Lee et al., 2020), faces (e.g. Rivolta et al., 2014), and auditory properties (e.g. Lee et al., 2011) among many other applications (Haynes & Rees, 2006).

Advances in the application of multivariate data driven methods to infant-friendly neuroimaging tools such as EEG and fNIRS bears promise for developmental researchers to begin answering questions beyond what can be addressed with traditional neuroimaging analysis techniques (Bayet et al., 2021; Norman et al., 2006; O’Brien et al., 2020; Zinszer et al., 2017). While this existing methodology lays the groundwork for infant study, challenges inherent to the collection and analysis of infant neural data require specific solutions and a thorough investigation into best practices. Infant data are often limited both by recruitment challenges, and extreme variation in usable trials due to unpredictable infant temperament, as well as movement when collecting neural data (Aslin & Fiser, 2005; Raschle et al., 2012). As a result, many methods of analyzing adult data may need to be modified to effectively implement them for infant research.

Recent work shows that applying MVPA to quantify the post-stimulus timecourse and representational characteristics of visual objects from infant EEG and fNIRS is feasible and opens new avenues for developmental research (Bayet et al., 2020; Emberson et al., 2017; Jessen et al., 2019; Mercure et al., 2020). In Bayet et al. (2020), EEG data from 12-15 month old infants as well as adults viewing images of animals and parts of the body were used to train a linear support vector machine (SVM) classifier, an analytic method that maps response features from neuroimaging data onto a high-dimensional space of stimulus dimensions. The accuracy of this stimulus-response mapping function is then assessed via 4-fold cross validation – a process of repeatedly training the SVM classifier on a subset of the data and testing that trained classifier on the withheld subset (25% = 4-fold). Patterns of activation in Bayet et al. yielded above-chance discrimination of 8 different visual stimuli in both adults and infants. Building on these results, here we outlined the steps required to perform time-resolved, within-subject MVPA with infant EEG data, summarized classification and validation best practices, and discussed the effectiveness of these methods given the limited number of trials typically available from infant EEG datasets. With this tutorial, we aim to make this powerful method more accessible, thereby expanding the tools available to developmental researchers.

## 2. Sample Dataset

The main idea of MVPA is to quantify the amount of information about a relevant dimension (e.g., was the stimulus a cat or a dog) that is available in the neural data, often by training and testing a classifier to discriminate between subsets of the neural data. As an example, here we focused on the pairwise decoding of visual stimuli from a sample dataset of infant (N=21) and adult (N=9) EEG data from a previously published report (Bayet et al., 2020). These data consisted of processed, normalized EEG voltages from 12-15-month-old infants and adults as they passively watched 8 static visual images of familiar animate objects (cat, dog, rabbit, bear, hand, foot, mouth, or nose). Processing steps that were applied to these EEG signals include line noise removal, filtering, re-referencing to the average, artifact removal, and epoching (Bayet et al., 2020). In addition, voltages were normalized by taking the z-score of the segmented EEG voltages with respect to the baseline period for each individual trial and channel (i.e., univariate noise normalization). The sample dataset is openly available at [Github link to be included after acceptance] as a .mat file.

## 3. MVPA Implementation

### 3.1 Programming implementations

To make EEG MVPA as widely accessible as possible, we provided a dual implementation of the core analysis code documented here in both MATLAB (R2019b) and Python (Python 3). Additional steps are provided in Python only. However, the libraries required have Matlab parallels, should one wish to implement them in Matlab as well. The clear advantage of Python is its portability and availability as an open source programming language. However, some matrix operations compute faster in Matlab. Both implementations produce comparable results, and a permutation based one-way ANOVA with cluster correction for multiple comparisons identified no clusters of difference within the data.

### 3.2 Cross-validation and pseudo-averaging

A key component of many MVPA implementations is the use of cross-validation. With cross-validation, only a portion of the available trials, the “training set”, is used to train the classifier. The remaining trials are held-out, forming the “test set”. A classifier is first trained on a substantial portion of the data from each participant (e.g., 75%) to estimate the activation patterns associated with the dimension or category of interest. Then the classifier’s performance is assessed based on its ability to use these estimates to make predictions about the withheld test set (**Figure 1**) (Bhavsar & Panchal, 2012). In this way, classification accuracy reflects the extent to which the classifier successfully extracted patterns from the training set that supported the discrimination of the relevant dimension in the training set (e.g., cat or dog) *and* that generalized to the test set. To avoid an idiosyncratic partitioning of the data into training and test sets, this procedure is repeated multiple times to randomly assign observations to the training and test sets. In our example, trial order was permuted (i.e., repeatedly sampled at random) within each participant and condition to form four folds (75%-25%) for cross-validation (Grootswagers et al., 2017). Previous work has demonstrated that such k-fold (here, k=4) cross validation techniques provide a more stable estimate of accuracy than comparable methods that have too many (such as leave-one-out) or too few (split-half) divisions of the entire dataset (Varoquaux et al., 2017).

**Figure 1.**
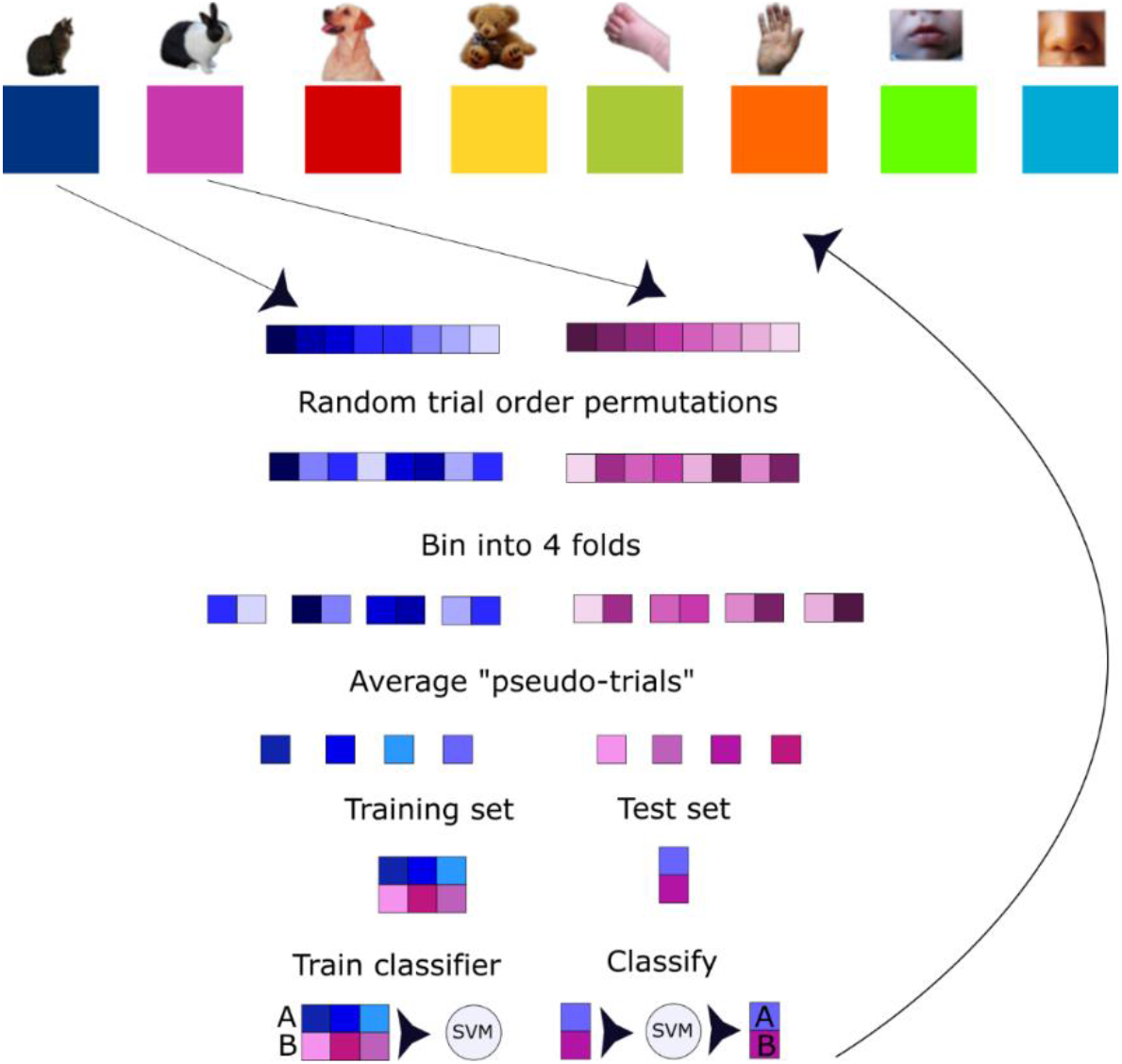
Process for pseudotrial generation and classification, repeated for all pairs of conditions. Available trials for each condition are randomly permuted, then divided into 4 bins of approximately equal size (+/-1 when trial number is not evenly divisible by 4). The trials in each bin are averaged to create 4 pseudotrials per condition, which are then used for training and testing the classifier. The resulting classification accuracies are averaged over all 200 trial order permutations for final pairwise accuracies.

The most straightforward way of implementing cross-validation is to treat each trial within a fold as an independent event. However, EEG data are noisy and so a more robust implementation involves some averaging of trials within each fold. For example, if there were two stimuli (e.g., cat or dog) and 20 repetitions of each with 4-fold cross validation, then within each fold the 5 trials of cat and the 5 trials of dog would be averaged, resulting in four “pseudotrials” (Grootswagers et al., 2017; Isik et al., 2014). Pairwise, within-subject classification of trials was performed such that two stimuli (e.g., cat vs. dog) were compared for each time point, with three pseudo-trials used for training and the fourth for testing. This procedure was repeated for 200 permutations of trial order, and classification accuracy was averaged over these permutations to yield a more robust estimate (Bayet et al., 2020) (**Figure 1**).

In some cases, additional testing of the model on an independent dataset can be desirable, going beyond cross-validation. For example, if researchers use cross-validation accuracy as a guide to choose their classification model (e.g., deciding on features, classifier type, or kernel based on which decision yields the highest cross-validation accuracy), then cross-validation alone would provide an overly optimistic estimate of the final model’s performance (Hastie et al., 2009). Even when it is not used to guide model selection, certain research questions may necessitate to assess model generalization beyond the parameters of a specific dataset (e.g., if assessing biomarkers, or if seeking to assess the generalizability of individual participants’ neural representations across multiple days, etc.). In such cases, testing the final model on additional held-out data may be required to better estimate the model’s performance.

### 3.3 Choosing response features to be used for classification

In this example, normalized voltage values across channels were used as features to train the classifier independently for each time point. The resulting decoding accuracy function represents how effectively the normalized amplitude values across channels predict which stimulus was present on a given trial in the test set at each time point after stimulus onset. Alternatively, researchers may wish to implement MVPA with summary statistics (e.g., average voltage during a rolling time-window) or spectral power across channels and frequency bands instead of voltages (Xie et al., 2020). Both feature approaches (i.e., time-domain and frequency-domain) have been demonstrated to effectively decode stimuli; however, at least in certain paradigms, different features may reflect different aspects of perception, cognition, or attention (Desantis et al., 2020). For example, Desantis et. al. demonstrated that voltage amplitude and alpha-band power both reliably decoded attention orientation, however alpha-band power was more associated with attention orienting in space while voltage amplitude signaled perceptual processes associated with attention. While the specific differences between frequency and amplitude measures of neural representations are unclear, both can be effective for decoding. In practice, using frequency features increases the granularity of the data because each channel provides multiple frequency-band components (e.g., alpha, theta, delta, gamma), thereby increasing signal to noise ratio. However, these frequency components must be extracted over a temporal window, thereby resulting in some loss of temporal resolution and increase in the potential dimensionality of the data (Vidaurre et al., 2020).

### 3.4 Choosing a classification algorithm

Here, we utilized a linear SVM to classify patterns of voltages across channels at each time-point. The tools leveraged for Matlab and Python were Libsvm (Chang & Lin, 2011) and scikit learn SVC (Pedregosa et al., 2011), respectively. The scikit learn SVC implementation is based on Libsvm and yields comparable results. Libsvm supports several variations to the SVM classifier. In the Python implementation all arguments to SVC were left as defaults. The Matlab implementation specifies a linear kernel, and a multi-class classification in the call to the SVM training function. The SVM classification method, which generates hyperplanes that maximize separation between categories in a high dimensional space, is particularly effective given the large number of features considered for classification in comparison to the small available number of training trials (observations) (Bhavsar & Panchal, 2012). SVM classifiers select samples that maximize the distance between categories, or support vectors to define the margins between categories. Support vectors are calculated such that they maximize the distance between the support vectors and the hyperplane that divides the categories. The decision boundaries defined in the training step are then used to classify the test data.

Alternatives to a linear SVM classifier include non-linear classifiers (e.g., Gaussian kernel SVM, Deep Neural Network) as well as other types of linear classifier such as logistic regression, Linear Discriminant Analysis, etc. Previous MVPA work suggests that most linear classification methods should perform similarly, as measured by prediction accuracy and stability of weights (Varoquaux et al., 2017). While a non-linear classifier can account for significantly more features than a linear approach, without a very large sample size such classification models are prone to overfitting (D’souza et al., 2020), i.e., fitting spurious patterns in the training data. It is also important to note that the SVM seeks <u>any</u> difference in the high-dimensional representation of the EEG features, including noise in the data. Since noise is a function of sample size, the success of the classifier could be due to a mismatch in the number of trials per stimulus rather than the underlying EEG features.

## 4. Resulting metrics and statistical testing

### 4.1 Output

In the provided pipeline, the output of the decoding function (in both language implementations) will be a Matlab (.mat) file containing the fields ‘out’ and ‘results’. The ‘out’ field contains the string name of the file. The ‘results’ field contains a 4-d double matrix of the resulting decoding accuracies ‘DA’, a structure containing the decoding parameters ‘params_decoding’, a matrix containing the number of trials completed for each participant in each condition ‘nreps’, and an array ‘times’ that is a list of all time points.

The ‘DA’ field is a 4-d matrix of the shape (number of participants, number of timepoints, number of conditions, number of conditions). That is, for each participant, at each timepoint, there is an upper diagonal matrix of average pairwise decoding accuracies for each stimulus pair. Of note, to avoid duplication, only the upper diagonal matrix (i.e., matrix elements above the diagonal) will contain numbers, while the diagonal and lower diagonal matrix will contain NaNs (not a number).

### 4.2 Within-subject pairwise classification accuracy against chance

To assess overall classification accuracy across the timeseries, the decoding accuracy (DA) matrix can be averaged over all subjects and conditions and compared to chance (50% in the case of pairwise classification). To derive an average timeseries over all participants, the condition by condition matrices need to be averaged over participants (i.e., the first, third, and fourth dimensions of the matrix in either Python or Matlab). This results in a one-dimensional array containing one average accuracy value per time point. To examine the pairwise decoding accuracy over the time series for each participant separately, accuracies should only be averaged over conditions (i.e., only the third and fourth dimensions): This results in a matrix of size (number of participants, number of time points) containing average classification accuracies at each time point for each participant. (**Figure 2**).

**Figure 2.**
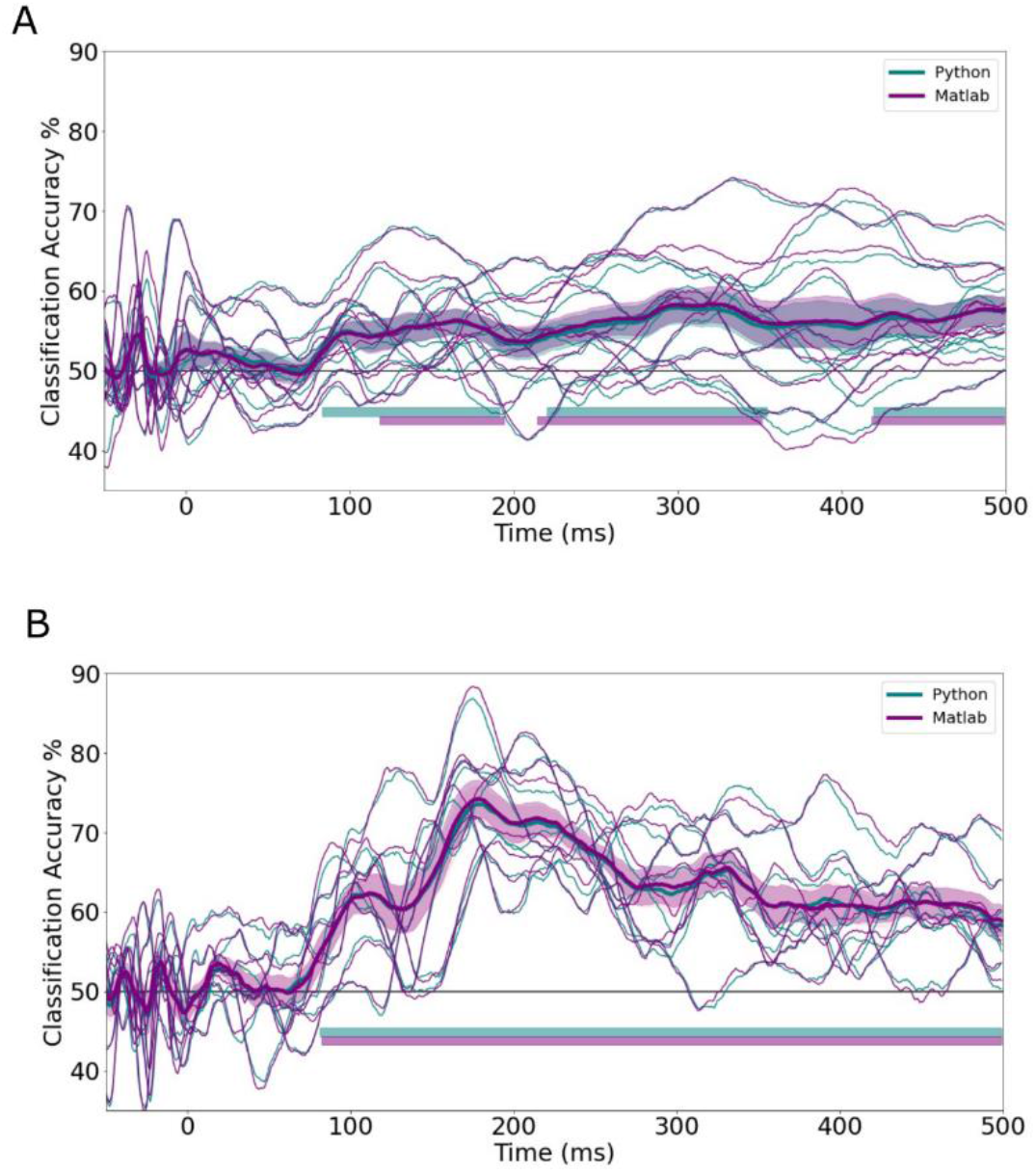
Mean overall decoding accuracy across the time series as generated by the Matlab and Python implementations for infants (**B**, n=10), and adults (**A**, n=8) with standard error highlighted. Time windows of cluster corrected above chance accuracy are denoted by the horizontal solid lines.

In our example, the significance of the classification accuracy against chance was calculated using a one-way right-tailed *t*-test, with cluster-based correction for multiple comparisons (Maris & Oostenveld, 2007). Alternatively, non-parametric equivalents (e.g., permutation-based) may be used. Of note, standard parametric or non-parametric statistical methods applied to classification accuracy do not support population level inference (Allefeld et al., 2016). In other words, because the actual value of the estimated classification accuracy can never be below chance, a *t*-test can only suggest that there is an effect in some individuals in the sample, a conclusion that does not generalize to the population. If population inference is necessary, alternative strategies have been proposed, such as examining the prevalence of the observed effect in the sample, as opposed to group means (Allefeld et al., 2016).

### 4.3 Representational Similarity Analyses

Representational similarity analysis (RSA) is a multivariate analysis method that assesses and compares the implied “geometry” of neural representations, i.e., how similar, or dissimilar patterns of neural activity are in response to different stimuli (Diedrichsen & Kriegeskorte, 2017; Haxby et al., 2014; Kriegeskorte & Kievit, 2013). The resulting measures of similarity or dissimilarity may then be compared between processing stages, groups, task conditions, or species, or between experimental and model data(Diedrichsen & Kriegeskorte, 2017; Haxby et al., 2014; Kriegeskorte & Kievit, 2013). In other words, RSA projects response differences from any dependent variable into a common space, thereby allowing those response differences to be compared with other responses differences or measures of difference regardless of the measures themselves (e.g., EEG, fMRI, model responses, behavioral ratings of dissimilarity) (Anderson et al., 2016; Bayet et al., 2020). Dissimilarity can be quantified in multiple ways such as within-class distance, Euclidean distance, pairwise correlations, and decoding accuracy (Guggenmos et al., 2018). Here we focused on classification accuracy, which is directly available from standard MVPA decoding, and cross-validated Euclidean distance, which has shown particular reliability as a measure of dissimilarity (Guggenmos et al., 2018).

The first step to RSA is constructing representational dissimilarity matrices (RDMs), which describe the difference between EEG feature patterns for the classes of stimuli (**Figure 3**). The accuracy based RDM is simply a matrix of pairwise classification accuracies across the set of stimuli. Measuring representational similarity based on Euclidean distance requires a separate decoding step. The procedure for Euclidean decoding was much the same as decoding with SVM, however the Euclidean distance between values, with additional cross-validation steps to improve signal-to-noise ratio, was calculated instead of classification accuracy. Following the formula described by Walther et. al., the difference between the mean EEG voltage values for two stimuli were calculated for test and training sets of pseudotrials, and multiplied (Walther et al., 2016). This created a more stable estimate of representational difference, given that noise is assumed to be independent between the two sets. Euclidean based RDMs were calculated using the same procedure described above. Regardless of how dissimilarity is calculated, decoding accuracy is based on which RDM from the training set is most similar to the RDM in the test set. RDMs can be used to test computational and cognitive theories, and allows for the comparison of representations without identifying the transformation between representational spaces (Kriegeskorte & Kievit, 2013). Note, in practice the ability to make group comparisons with RSA is limited by the number of stimuli. In this example, RDMs contain 28 distances between pairs of 8 stimuli; based on this number of distances, analyses correlating RDMs between groups or time-windows can theoretically detect correlations of *r*∼0.45 or higher with 80% power (one-tail linear correlation, α = 5% for a single test; G*Power 3.1).

**Figure 3.**
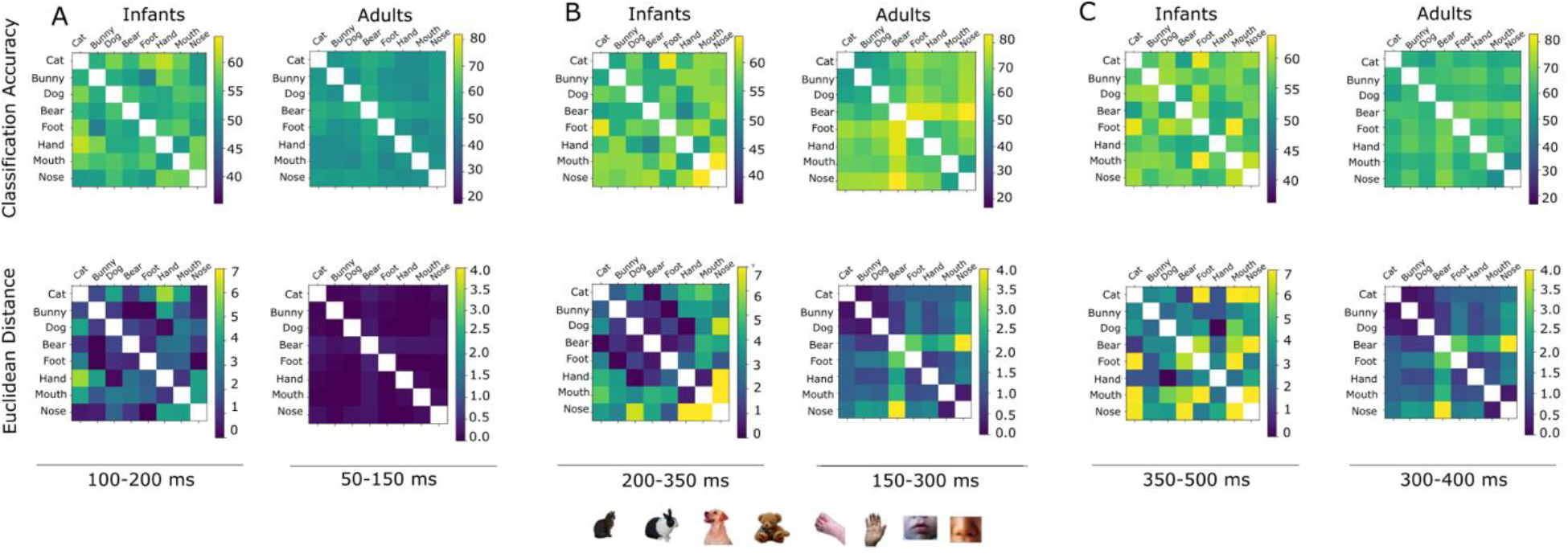
Representational dissimilarity matrices of pairwise classification accuracy and cross validated Euclidean distance for the subsets of adults (n=8) and infants (n=15) with highest overall RDM reliability. RDMs calculated in the time windows during which classification accuracy rises above chance (**A**), during the window of highest classification accuracy (**B**) and following the window of highest classification accuracy (**C**).

## 5. Impact of limited trial numbers and criteria for participant inclusion

Difficulties collecting enough valid trials for analysis frequently impede infant research. A range of valid trials thresholds were tested here to assess the relative impact of the number of available valid trials on the stability of decoding accuracy within the test set. At each of the trial thresholds, decoding was performed on all participant data with enough artifact-free trials, both cut off at exactly the trial threshold (i.e., at a threshold of 4, 4 trials from each condition were randomly selected for analysis if a participant had more than 4 available) and including all available trials. Within the example datasets, the number of participants included in the analysis was reduced as the trial threshold for participant inclusion in the analysis became more stringent (**Figure 4**).

**Figure 4.**
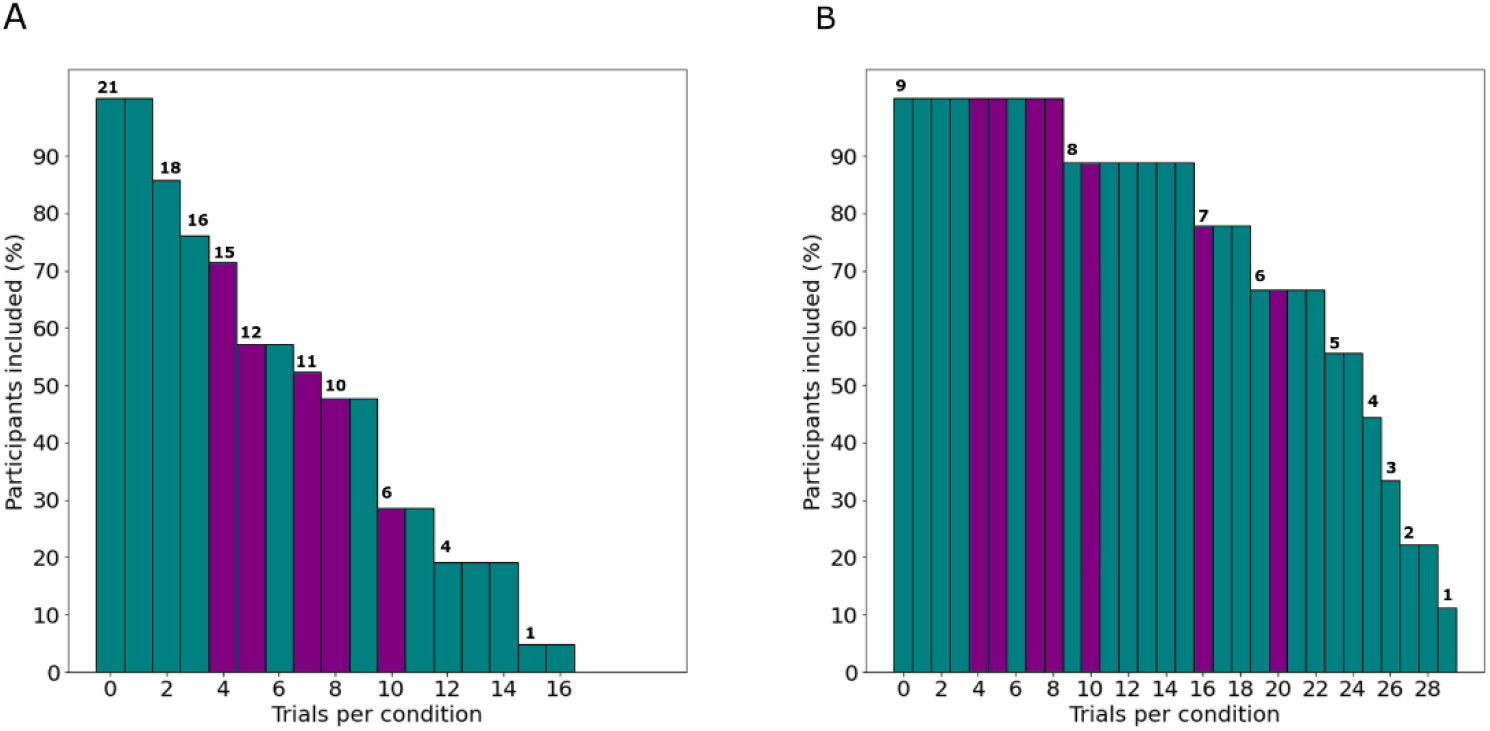
Number of participants from the test data included vs. trial threshold for infants (**A**) and adults (**B**). Trial thresholds tested are highlighted in purple, and number of participants included at each threshold are noted at the top of the bars.

### 5.1 Impact on classification accuracy

In both infants and adults there was an expected decrease in decoding accuracy when trials were cut off at the threshold compared to when all available trials were used. The results showed very similar time windows of above chance accuracy regardless of trial number threshold in both the infant and adult data (**Figure 5**). Higher numbers of valid trials led to higher classification accuracy in the adult dataset as expected. However, this pattern was perhaps less marked in the infant dataset (**Figures 5-6**), presumably due to a ceiling effect as well as to some level of trade-off between the number of available trials and the number of available participants with at least that number of available trials. Decoding accuracy in infants was numerically higher at the most stringent threshold of 10 trials per condition, however this pattern of results may reflect the particularities of the small amount of participant data included (**Figure 6**). In general, it is not possible to state a priori how many valid trials per stimulus are required to generate asymptotic decoding accuracy because of differences in the discriminability among a set of stimuli. However, analysis of reliability tradeoff when lowering the trial threshold in pilot data may provide guidance on this issue.

**Figure 5.**
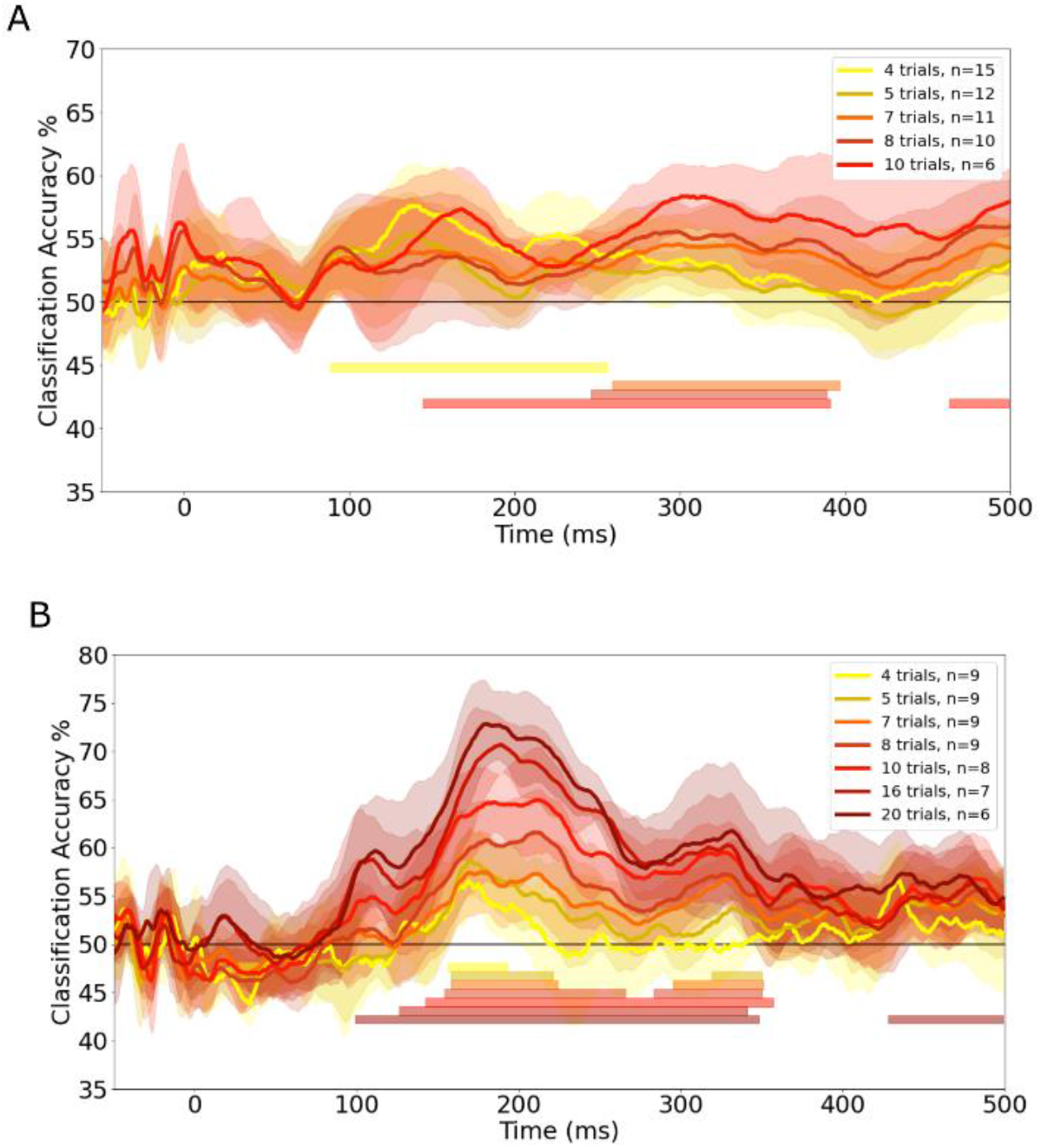
Overall average decoding accuracy when number of trials per condition was restricted to exactly each of the trial thresholds, with 95% confidence interval highlighted, for (**A**) infants and (**B**) adults. Time windows of cluster corrected above chance accuracy are denoted by the horizontal solid lines. Participants with fewer than the specified number of artifact-free trials are excluded (see Figure 1).

**Figure 6.**
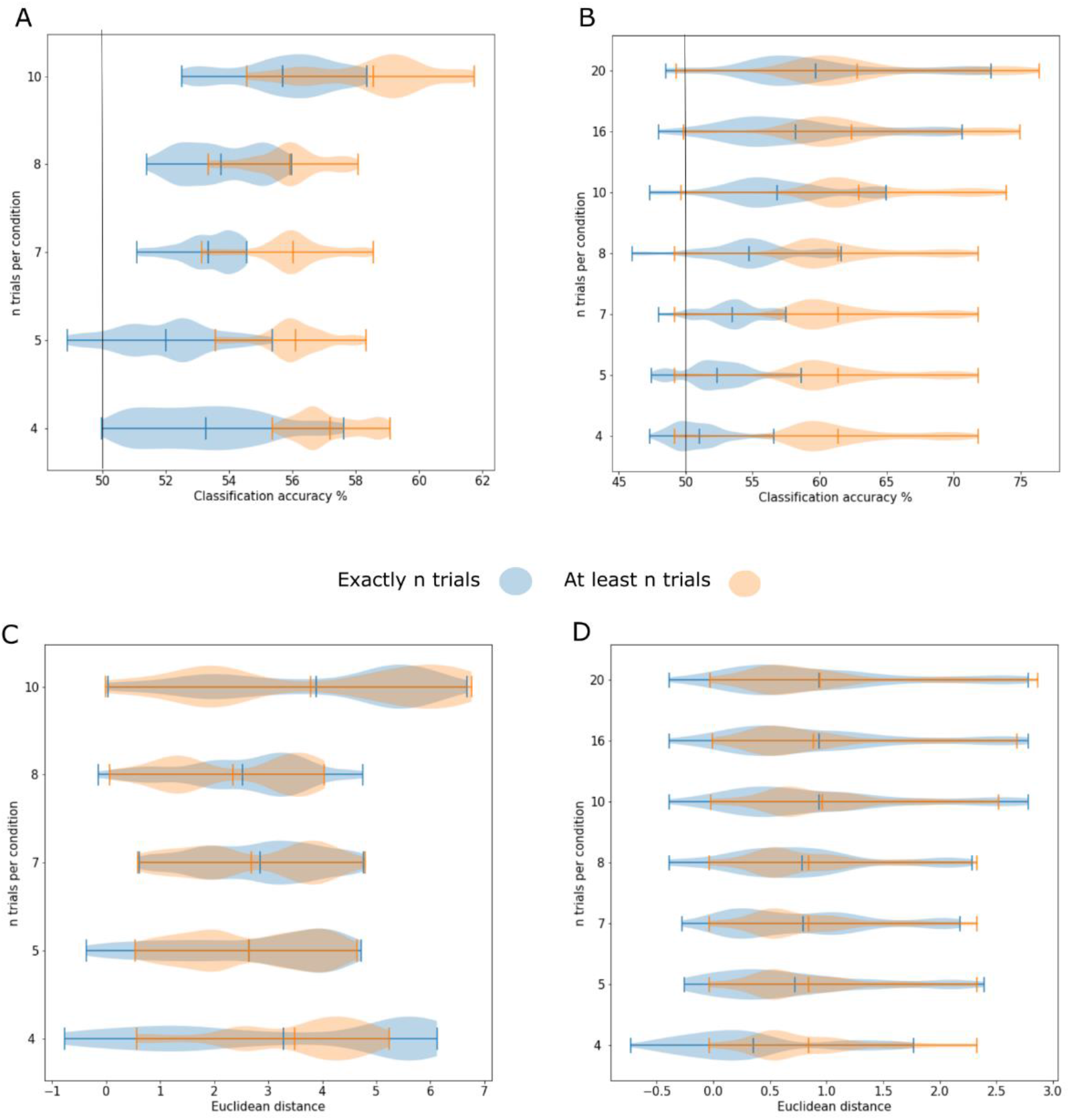
Average classification accuracy at different trial thresholds with (**A**) infants (time window 100-500 ms) and (**B**) adults (time window 50-500 ms) and Euclidean distance with (**C**) infants (time window 100-500 ms) and (**D**) adults (time window 50-500 ms). Blue denotes the distribution when the number of trials included was cut off at the threshold, and orange denotes when all trials were included for all participants who met the threshold of trials per condition.

### 5.2 Impact on the reliability of Representational Dissimilarity Matrices

To assess the feasibility of RSA in the infant EEG dataset, we also examined the reliability by which the dissimilarity between neural responses to different stimulus types could be estimated at the group level – i.e., the noise ceiling (Nili et al., 2014). To that end, we used the Spearman-Brown split-half reliability method which involves correlating dissimilarity matrices, composed of the pairwise dissimilarities between all stimuli pairs, between two halves of the dataset (Lage-Castellanos et al., 2019). Specifically, the Pearson’s correlation coefficient was calculated between group-level RDMs estimated from random half-splits of the full group, repeated for 100 split halves (Nili et al., 2014). The statistical significance of these estimates was determined by repeating the same split half procedure, only this time shuffling dissimilarities in one of the splits in each iteration, repeated for 10,000 permutations (Lage-Castellanos et al., 2019) to form a null distribution against which to compare empirical reliability values (**Figure 7**).

**Figure 7.**
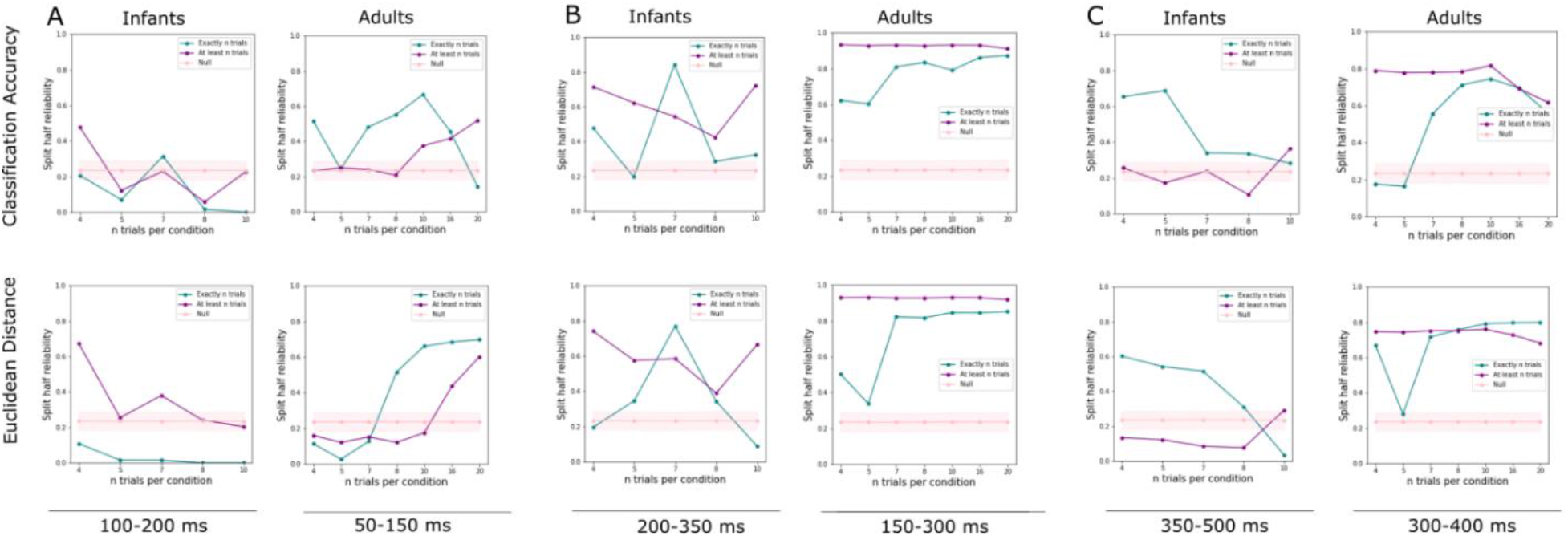
Average pairwise split-half reliability of the group-level Representational Dissimilarity Matrices of both classification accuracy and Euclidean distance obtained at each trial number threshold with corresponding average and 2.5-97.5 percentiles of the null split-half noise ceiling calculated in the time windows preceding above chance classification (**A**), during the window of highest classification accuracy (**B**) and following the window of highest classification accuracy (**C**).

In the adult dataset, the number of artifact-free trials used as inclusion threshold was numerically correlated with the observed split-half reliability of the corresponding group-level RDMs when including exactly N trials per condition in decoding, i.e., there was a trend for reliability to decrease when decreasing the amount of available data. When decoding was performed using all available data in adults who met the trial threshold for inclusion, there was little change between thresholds and an overall negative correlation between trial threshold. The pattern was less clear in infants, where numerically the number of artifact free trials used as inclusion threshold was negatively correlated with reliability when including exactly n trials per condition in decoding, and when using all available data from included participants (**Figure 7**). Correlations were not statistically significant but suggest that, for group-level RSA with small infant datasets, decreasing the number of artifact-free trials needed for participant-inclusion may not necessarily decrease how reliable the resulting group RDMs are, and may in fact yield more reliable estimates if the number of included participants can be increased (i.e., there is a trade-off between the number of trials per stimulus per infant and the number of infants).

## 6. Discussion

This tutorial aims to expand access to time-resolved MVPA and facilitate its future application to novel developmental research. Due to the number of logistical difficulties involved with collecting fMRI data from awake infants (for examples where this is successfully done, see e.g. (Dehaene-Lambertz et al., 2002; Ellis & Turk-Browne, 2018) and the relative ease of collecting EEG data, a standard methodology for applying MVPA to infant EEG is extremely valuable. Providing implementations in two commonly used programming languages (Matlab, Python) significantly increases the availability of this method. As demonstrated here, both implementations give comparable results. Both infant and adult EEG data were successfully used to achieve reliable decoding of two or more stimuli, with infant classification at above chance levels even with restrictions on trial numbers.

There are several important limitations to MVPA as a means of accessing neural representations. First, similar to univariate analyses, MVPA is sensitive to any pattern that differentiates categories. It is not guaranteed that the underlying cause of such a multivariate pattern is a cognitive process of interest, as opposed to some spurious factor such a low-level difference in stimulus brightness, size, or number of trials. Second, while linear classification requires fewer data to yield robust results than non-linear methods such as artificial neural networks (Alwosheel et al., 2018), this method is limited by the assumption of linearity inherent in the classification method. While there are theoretical and practical reasons to favor the use of linear classifiers when employing MVPA to assess neural representations (Hung et al., 2005; King et al., 2018), there is always a possibility that discrimination within the brain relies on nonlinear patterns of activation that do not fall within the linear constraints of the classifier (Naselaris & Kay, 2015; Popov et al., 2018).

There are also caveats to the current example results that should be kept in mind when implementing this method. Primarily, the data set used to produce example decoding results was small. While the method was successfully executed on these data, the limited sample size could have skewed the presented results. It is also worth noting that there could be discrepancies that would be apparent with a larger data set. These specific findings also may not generalize to other sensory domains, EEG sensor types, or age groups. In a similar vein, the example results shown here may not generalize to other kinds of visual stimuli. Future research may address these limitations by replicating the current analyses in different, larger datasets of infant EEG data.

By applying MVPA to infant EEG data in a pre-verbal age group, developmental researchers can draw conclusions about the nature and consistency of neural representations of perceived stimuli beyond what is afforded by univariate behavioral or neuroimaging methods. Future research may further expand the use of MVPA with infant data to other neuroimaging modalities (e.g., fMRI, time-frequency decomposition of EEG data, source-localized EEG data) and tailor data collection and analysis methods to better address limitations of infant neuroimaging including the quality and quantity of data available for data-intensive analyses such as MVPA.

## Acknowledgments

We thank the participants, families, research assistants, and students who made this work possible.

## References

Allefeld, C., Görgen, K., & Haynes, J.-D. (2016). Valid population inference for information-based imaging: From the second-level t-test to prevalence inference. NeuroImage, 141, 378–392. https://doi.org/10.1016/j.neuroimage.2016.07.040

Alwosheel, A., van Cranenburgh, S., & Chorus, C. G. (2018). Is your dataset big enough? Sample size requirements when using artificial neural networks for discrete choice analysis. Journal of Choice Modelling, 28, 167–182. https://doi.org/10.1016/j.jocm.2018.07.002

Anderson, A. J., Zinszer, B. D., & Raizada, R. D. S. (2016). Representational similarity encoding for fMRI: Pattern-based synthesis to predict brain activity using stimulus-model-similarities. NeuroImage, 128, 44–53. https://doi.org/10.1016/j.neuroimage.2015.12.035

Aslin, R. N., & Fiser, J. (2005). Methodological challenges for understanding cognitive development in infants. Trends in Cognitive Sciences, 9(3), 92–98. https://doi.org/10.1016/j.tics.2005.01.003

Bayet, L., Saville, A., & Balas, B. (2021). Sensitivity to face animacy and inversion in childhood: Evidence from EEG data. Neuropsychologia, 156, 107838. https://doi.org/10.1016/j.neuropsychologia.2021.107838

Bayet, L., Zinszer, B. D., Reilly, E., Cataldo, J. K., Pruitt, Z., Cichy, R. M., Nelson, C. A., & Aslin, R. N. (2020). Temporal dynamics of visual representations in the infant brain. Developmental Cognitive Neuroscience, 45, 100860. https://doi.org/10.1016/j.dcn.2020.100860

Bell, M. A., & Cuevas, K. (2012). Using EEG to Study Cognitive Development: Issues and Practices. Journal of Cognition and Development, 13(3), 281–294. https://doi.org/10.1080/15248372.2012.691143

Bhavsar, H., & Panchal, M. H. (2012). A review on support vector machine for data classification. Int. J. Adv. Res. Comput. Eng. Technol, 185–189.

Brodersen, K. H., Wiech, K., Lomakina, E. I., Lin, C., Buhmann, J. M., Bingel, U., Ploner, M., Stephan, K. E., & Tracey, I. (2012). Decoding the perception of pain from fMRI using multivariate pattern analysis. NeuroImage, 63(3), 1162–1170. https://doi.org/10.1016/j.neuroimage.2012.08.035

Chang, C.-C., & Lin, C.-J. (2011). LIBSVM: A library for support vector machines. ACM Transactions on Intelligent Systems and Technology, 2(3), 27:1-27:27. https://doi.org/10.1145/1961189.1961199

Dehaene-Lambertz, G., Dehaene, S., & Hertz-Pannier, L. (2002). Functional Neuroimaging of Speech Perception in Infants. Science, 298(5600), 2013–2015. https://doi.org/10.1126/science.1077066

Dehaene-Lambertz, G., & Spelke, E. S. (2015). The Infancy of the Human Brain. Neuron, 88(1), 93–109. https://doi.org/10.1016/j.neuron.2015.09.026

Desantis, A., Chan-Hon-Tong, A., Collins, T., Hogendoorn, H., & Cavanagh, P. (2020). Decoding the Temporal Dynamics of Covert Spatial Attention Using Multivariate EEG Analysis: Contributions of Raw Amplitude and Alpha Power. Frontiers in Human Neuroscience, 14. https://doi.org/10.3389/fnhum.2020.570419

Diedrichsen, J., & Kriegeskorte, N. (2017). Representational models: A common framework for understanding encoding, pattern-component, and representational-similarity analysis. PLOS Computational Biology, 13(4), e1005508. https://doi.org/10.1371/journal.pcbi.1005508

D’souza, R. N., Huang, P.-Y., & Yeh, F.-C. (2020). Structural Analysis and Optimization of Convolutional Neural Networks with a Small Sample Size. Scientific Reports, 10(1), 834. https://doi.org/10.1038/s41598-020-57866-2

Ellis, C. T., & Turk-Browne, N. B. (2018). Infant fMRI: A Model System for Cognitive Neuroscience. Trends in Cognitive Sciences, 22(5), 375–387. https://doi.org/10.1016/j.tics.2018.01.005

Emberson, L. L., Zinszer, B. D., Raizada, R. D. S., & Aslin, R. N. (2017). Decoding the infant mind: Multivariate pattern analysis (MVPA) using fNIRS. PLOS ONE, 12(4), e0172500. https://doi.org/10.1371/journal.pone.0172500

Grootswagers, T., Wardle, S. G., & Carlson, T. A. (2017). Decoding Dynamic Brain Patterns from Evoked Responses: A Tutorial on Multivariate Pattern Analysis Applied to Time Series Neuroimaging Data. Journal of Cognitive Neuroscience, 29(4), 677–697. https://doi.org/10.1162/jocn_a_01068

Guggenmos, M., Sterzer, P., & Cichy, R. M. (2018). Multivariate pattern analysis for MEG: A comparison of dissimilarity measures. NeuroImage, 173, 434–447. https://doi.org/10.1016/j.neuroimage.2018.02.044

Hastie, T., Tibshirani, R., & Friedman, J. (2009). Linear Methods for Classification. In T. Hastie, R. Tibshirani, & J. Friedman (Eds.), The Elements of Statistical Learning: Data Mining, Inference, and Prediction (pp. 101–137). Springer. https://doi.org/10.1007/978-0-387-84858-7_4

Haxby, J. V. (2012). Multivariate pattern analysis of fMRI: The early beginnings. NeuroImage, 62(2), 852–855. https://doi.org/10.1016/j.neuroimage.2012.03.016

Haxby, J. V., Connolly, A. C., & Guntupalli, J. S. (2014). Decoding Neural Representational Spaces Using Multivariate Pattern Analysis. Annual Review of Neuroscience, 37(1), 435–456. https://doi.org/10.1146/annurev-neuro-062012-170325

Haynes, J.-D., & Rees, G. (2006). Decoding mental states from brain activity in humans. Nature Reviews Neuroscience, 7(7), 523–534. https://doi.org/10.1038/nrn1931

Hung, C. P., Kreiman, G., Poggio, T., & DiCarlo, J. J. (2005). Fast Readout of Object Identity from Macaque Inferior Temporal Cortex. Science, 310(5749), 863–866. https://doi.org/10.1126/science.1117593

Isik, L., Meyers, E. M., Leibo, J. Z., & Poggio, T. (2014). The dynamics of invariant object recognition in the human visual system. Journal of Neurophysiology, 111(1), 91–102. https://doi.org/10.1152/jn.00394.2013

Jessen, S., Fiedler, L., Münte, T. F., & Obleser, J. (2019). Quantifying the individual auditory and visual brain response in 7-month-old infants watching a brief cartoon movie. NeuroImage, 202, 116060. https://doi.org/10.1016/j.neuroimage.2019.116060

King, J.-R., Gwilliams, L., Holdgraf, C., Sassenhagen, J., Barachant, A., Engemann, D., Larson, E., & Gramfort, A. (2018). Encoding and Decoding Neuronal Dynamics: Methodological Framework to Uncover the Algorithms of Cognition. https://hal.archives-ouvertes.fr/hal-01848442

Kriegeskorte, N., & Kievit, R. A. (2013). Representational geometry: Integrating cognition, computation, and the brain. Trends in Cognitive Sciences, 17(8), 401–412. https://doi.org/10.1016/j.tics.2013.06.007

Lage-Castellanos, A., Valente, G., Formisano, E., & Martino, F. D. (2019). Methods for computing the maximum performance of computational models of fMRI responses. PLOS Computational Biology, 15(3), e1006397. https://doi.org/10.1371/journal.pcbi.1006397

Lee, I.-S., Jung, W., Park, H.-J., & Chae, Y. (2020). Spatial Information of Somatosensory Stimuli in the Brain: Multivariate Pattern Analysis of Functional Magnetic Resonance Imaging Data. Neural Plasticity, 2020, e8307580. https://doi.org/10.1155/2020/8307580

Lee, Y.-S., Janata, P., Frost, C., Hanke, M., & Granger, R. (2011). Investigation of melodic contour processing in the brain using multivariate pattern-based fMRI. NeuroImage, 57(1), 293–300. https://doi.org/10.1016/j.neuroimage.2011.02.006

Maris, E., & Oostenveld, R. (2007). Nonparametric statistical testing of EEG- and MEG-data. Journal of Neuroscience Methods, 164(1), 177–190. https://doi.org/10.1016/j.jneumeth.2007.03.024

Mercure, E., Evans, S., Pirazzoli, L., Goldberg, L., Bowden-Howl, H., Coulson-Thaker, K., Beedie, I., Lloyd-Fox, S., Johnson, M. H., & MacSweeney, M. (2020). Language Experience Impacts Brain Activation for Spoken and Signed Language in Infancy: Insights From Unimodal and Bimodal Bilinguals. Neurobiology of Language, 1(1), 9–32. https://doi.org/10.1162/nol_a_00001

Naselaris, T., & Kay, K. N. (2015). Resolving ambiguities of MVPA using explicit models of representation. Trends in Cognitive Sciences, 19(10), 551–554. https://doi.org/10.1016/j.tics.2015.07.005

Nili, H., Wingfield, C., Walther, A., Su, L., Marslen-Wilson, W., & Kriegeskorte, N. (2014). A Toolbox for Representational Similarity Analysis. PLOS Computational Biology, 10(4), e1003553. https://doi.org/10.1371/journal.pcbi.1003553

Norman, K. A., Polyn, S. M., Detre, G. J., & Haxby, J. V. (2006). Beyond mind-reading: Multi-voxel pattern analysis of fMRI data. Trends in Cognitive Sciences, 10(9), 424–430. https://doi.org/10.1016/j.tics.2006.07.005

O’Brien, A. M., Bayet, L., Riley, K., Nelson, C. A., Sahin, M., & Modi, M. E. (2020). Auditory Processing of Speech and Tones in Children With Tuberous Sclerosis Complex. Frontiers in Integrative Neuroscience, 14. https://doi.org/10.3389/fnint.2020.00014

Pedregosa, F., Varoquaux, G., Gramfort, A., Michel, V., Thirion, B., Grisel, O., Blondel, M., Prettenhofer, P., Weiss, R., Dubourg, V., Vanderplas, J., Passos, A., & Cournapeau, D. (2011). Scikitlearn: Machine Learning in Python. Journal of Machine Learning Research, 12, 2825–2830.

Popov, V., Ostarek, M., & Tenison, C. (2018). Practices and pitfalls in inferring neural representations. NeuroImage, 174, 340–351. https://doi.org/10.1016/j.neuroimage.2018.03.041

Raschle, N., Zuk, J., Ortiz-Mantilla, S., Sliva, D. D., Franceschi, A., Grant, P. E., Benasich, A. A., & Gaab, N. (2012). Pediatric neuroimaging in early childhood and infancy: Challenges and practical guidelines. Annals of the New York Academy of Sciences, 1252, 43–50. https://doi.org/10.1111/j.1749-6632.2012.06457.x

Rivolta, D., Woolgar, A., Palermo, R., Butko, M., Schmalzl, L., & Williams, M. A. (2014). Multi-voxel pattern analysis (MVPA) reveals abnormal fMRI activity in both the “core” and “extended” face network in congenital prosopagnosia. Frontiers in Human Neuroscience, 8. https://doi.org/10.3389/fnhum.2014.00925

Varoquaux, G., Raamana, P. R., Engemann, D. A., Hoyos-Idrobo, A., Schwartz, Y., & Thirion, B. (2017). Assessing and tuning brain decoders: Cross-validation, caveats, and guidelines. NeuroImage, 145, 166–179. https://doi.org/10.1016/j.neuroimage.2016.10.038

Vidaurre, D., Cichy, R. M., & Woolrich, M. W. (2020). Dissociable components of oscillatory activity underly information encoding in human perception. BioRxiv, 2020.09.10.291294. https://doi.org/10.1101/2020.09.10.291294

Walther, A., Nili, H., Ejaz, N., Alink, A., Kriegeskorte, N., & Diedrichsen, J. (2016). Reliability of dissimilarity measures for multi-voxel pattern analysis. NeuroImage, 137, 188–200. https://doi.org/10.1016/j.neuroimage.2015.12.012

Xie, S., Kaiser, D., & Cichy, R. M. (2020). Visual Imagery and Perception Share Neural Representations in the Alpha Frequency Band. Current Biology, 30(13), 2621-2627.e5. https://doi.org/10.1016/j.cub.2020.04.074

Zinszer, B. D., Bayet, L., Emberson, L. L., Raizada, R. D. S., & Aslin, R. N. (2017). Decoding semantic representations from functional near-infrared spectroscopy signals. Neurophotonics, 5(1), 011003. https://doi.org/10.1117/1.NPh.5.1.011003

